# Micro-Scale Transport Processes for Advanced Healthcare and Point of Care Diagnostics

**DOI:** 10.1101/113787

**Authors:** Aashish Priye

## Abstract

Recent outbreaks like Zika and Ebola highlights the challenges associated with pathogen diagnostics in the developing world. With the outbreak in Africa, and isolated cases on other continents, the need for an affordable, rapid and portable diagnostic solution has been repeatedly stressed and is one of the most critical issues confronting global health. Unfortunately, the current conventional PCR instrumentation needed to perform gold standard DNA-based diagnostic tests is bulky, slow, and expensive, making it unsuitable for resource-limited settings in developing countries where dedicated laboratory facilities are not available. Advances in micro-fluidics and smartphone based technology has paved way for novel implementations of traditional molecular diagnostic techniques. We present a series of advances in molecular diagnostic techniques by harnessing convective flow to actuate biochemical reactions.

## INTRODUCTION

Recently, it has been shown that convective flow fields can be harnessed to perform PCR. Our research led us to investigate the underlying physics of these thermally driven micro-scale Rayleigh-Bénard convective flows. This new understanding revealed how a subset of these flow fields exhibit chaotic advection that greatly accelerate temperature driven biochemical reactions such as PCR [1–6]. Since its first introduction a little over a decade ago, convective thermo-cycling has remained an intriguing avenue to enable rapid PCR. However, a crucial roadblock to practical implementation of this approach has been the inherent interdependence between the internal flow field and the reactor geometry (expressed in terms of the height, h, and diameter, d, of a cylindrically shaped configuration). The spatial temperature gradient established between the top (cool) and bottom (hot) surfaces of the reactor not only actuates the denaturing, annealing, and extension steps necessary to perform the PCR, it also supplies the driving force to physically transport reagents between these reaction zones. It has previously been assumed that this interplay implies a need to custom design reactor geometries to match the individual thermal requirements of each PCR assay to be performed (e.g., when different primers with different annealing temperatures are employed), and that robustness is constrained by a need to maintain a specific orientation with respect to the gravitational driving force.

**Figure 1:**
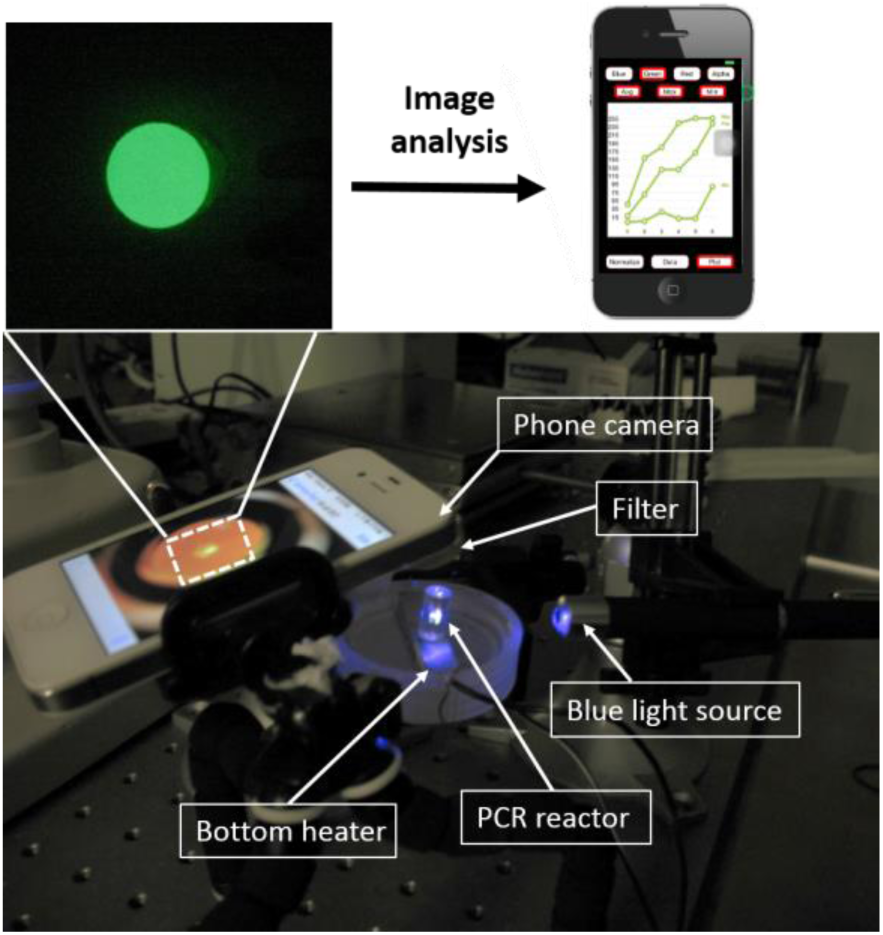
Real time fluorescent detection of convective PCR products via SYBR-PCR chemistry. A blue light source illuminates the convective PCR reactor and the images of the reactor top are taken with smart-phone camera which are subsequently analyzed with a built in image analysis app called PCRtoGo.

## PORTABLE PCR SETUP

We recently developed a 3D coupled flow-reaction model that reveals an unexpectedly broad design space dominated by chaotic advection where reaction rates are greatly accelerated and remain essentially unchanged over virtually the entire range of realistic PCR condition. Any reactor geometry selected within this regime is therefore universally functional (i.e., analogous to standardized PCR tubes and plates), making it possible to execute a 30 cycle PCR in 10 – 20 min regardless of temperature setpoints and spatial orientation. Having found the optimal design parameters for convective PCR, we coupled this novel isothermal setup with a versatile smartphone based detection unit and an integrated image analysis app [7–9]. This greatly simplifies the design, enabling us to integrate such biochemical analysis platforms with consumer-class quadcopter drones for rapid deployment of nucleic acid based diagnostics [10]. The ability to perform rapid in-flight assays with smartphone connectivity eliminates delays between sample collection and analysis so that test results can be delivered in minutes, suggesting new possibilities for drone-based systems to function in broader and more sophisticated roles beyond cargo transport and imagine.

**Figure 2.**
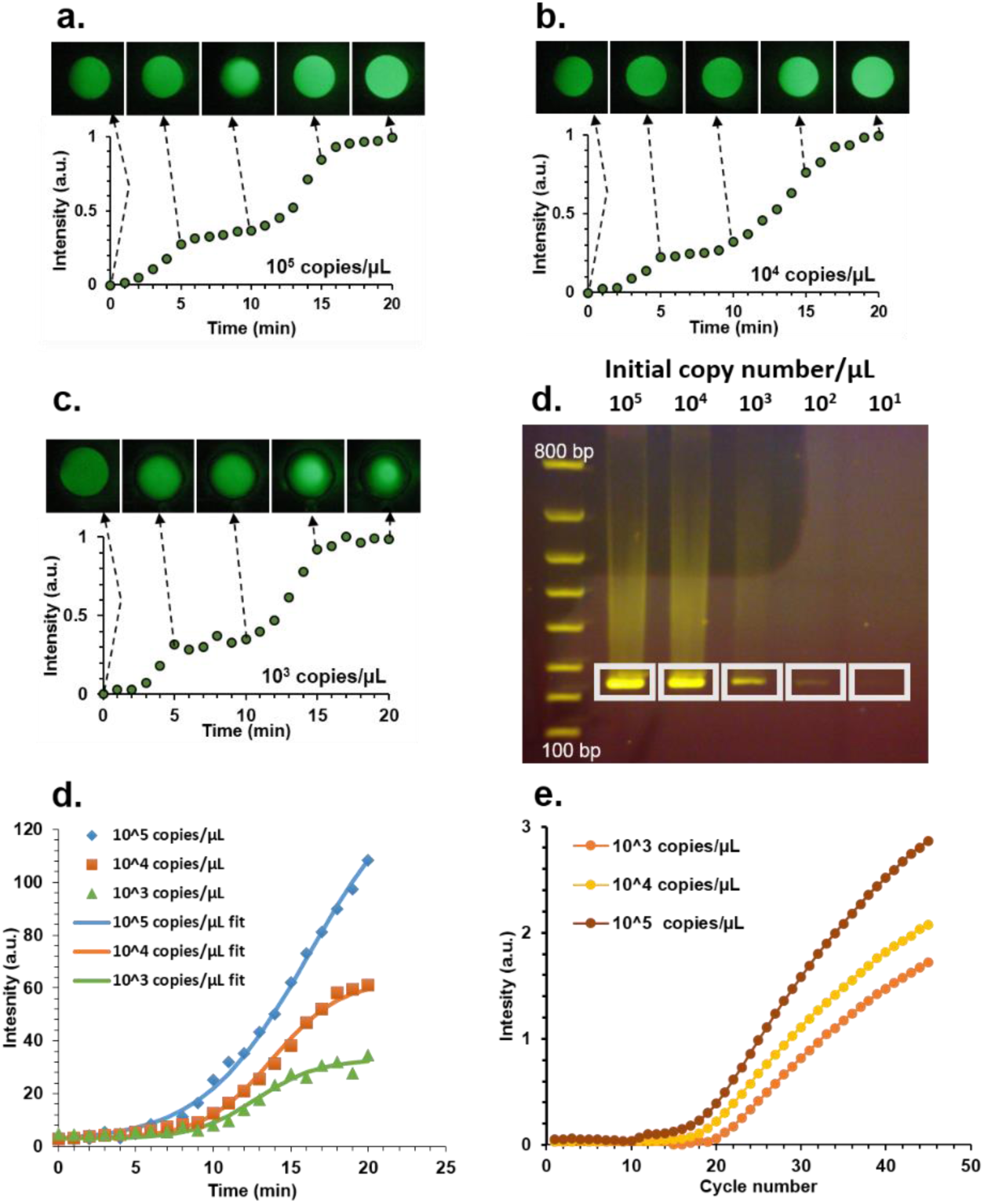
Smart-phone based PCR detection. (a, b, c) Real time quantification of a targeted 237 bp segment of lambda DNA for 10^5^, 10^4^ and 10^3^ initial DNA copy numbers respectively. (d) Gel electropherograms revealed that the convective PCR is capable of amplifying the targeted sequence from an initial copy number as low as ~100 copies/uL. (e) Average real time quantification using PCRtoGo app with gamma correction and corresponding sigmoidal curve fit. (f) Real time PCR run on a benchtop qPCR machine (Roche).

## CONVECTIVE FLOWS AND SURFACE CHEMISTRY

The surprising interplay between reactions and micro-scale convective flows led us to consider adaptations beyond PCR. Specifically, we demonstrate that such flows, naturally established over a broad range of hydrothermally relevant pore sizes, function as highly efficient conveyors to continually shuttle molecular precursors from the bulk fluid to targeted locations on the solid boundaries, enabling greatly accelerated chemical synthesis. These porous mineral formations embed richly complex microenvironments capable of catalytically polymerizing monomers and orchestrating fundamental electrochemistry central to prebiotic evolution of metabolic processes. But a unified framework explaining how surface-mediated synthesis can be orchestrated by the interplay among physical, chemical, and thermal processes within these catalytically active networks remains elusive. We explored the emerging convective flows in hydrothermally relevant pore sizes and discovered that they have the capacity to act as highly efficient conveyors to continually shuttle molecular precursors from the bulk fluid to targeted locations on the solid boundaries where they assemble into membrane-like films capable of electrochemically generating pH gradients. We quantitatively mapped the enrichment of biomolecular species achievable via this process, and introduced an in situ approach to directly probe its influence on surface reaction kinetics. Our results suggest that chaotic thermal convection may supply a previously unappreciated driving force to support emergence of early bioenergetic pathways—a key unanswered question in the origin of life [11–15].

**Figure 3:**
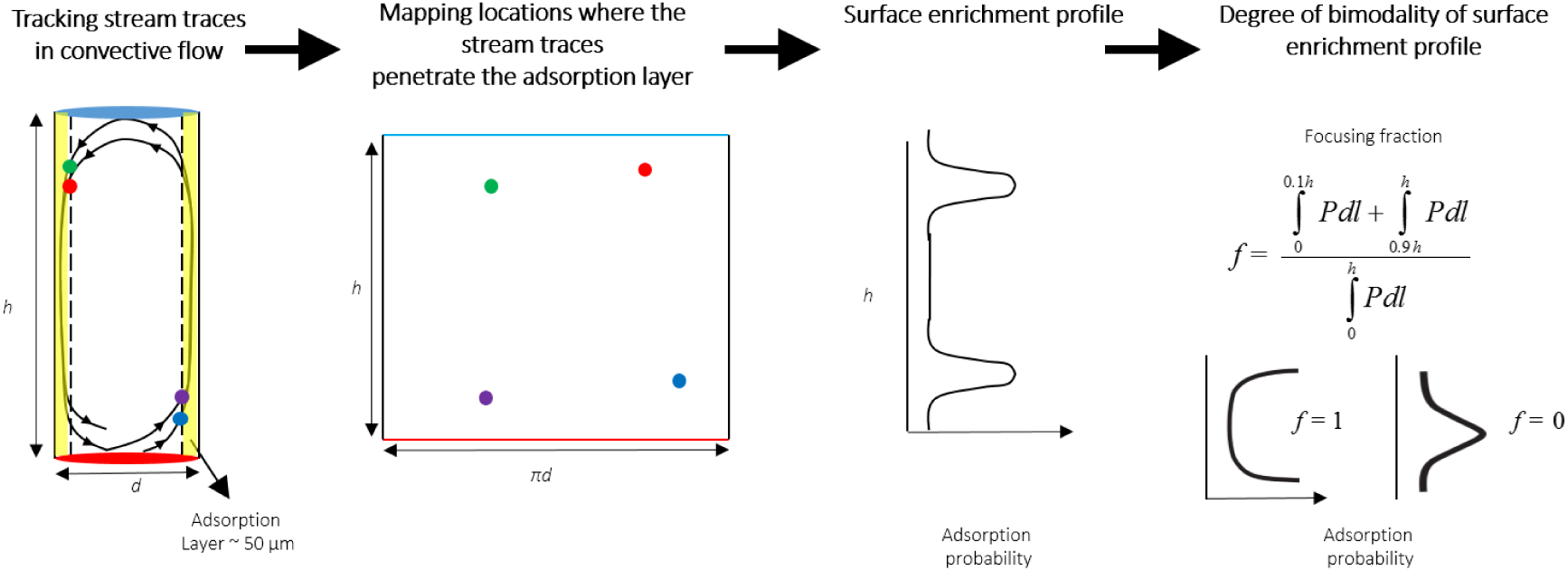
Construction of pore boundary adsorption profile. The fluid elements are tracked as they are converted within the pore and the locations where the stream traces penetrate the boundary layer are mapped. Subsequently an adsorption profile is constructed to quantify the focusing fraction.

## MICROSCALE ADHESION MODEL

Another aspect of my research involves the hydro dynamical interaction between micron-sized particles in intricate flow fields. Our computational fluid dynamics models have thrown light on how the microluidic flow fields can be tune fluid-particle interactions [13]. We development of a new adhesion model for particle resuspension modeling. The phenomenon of particle resuspension plays a vital role in numerous fields. Among many aspects of particle resuspension dynamics, a dominant concern is the accurate description and formulation of the van der Waals (vdW) interactions between the particle and substrate. Current models treat adhesion by incorporating a material dependent Hamaker's constant which relies on the heuristic Hamaker's two body interactions.

However, this assumption of pair wise summation of interaction energies can lead to significant errors in condensed matter as it does not take into account the many body interaction and retardation effects. To address these issues, an approach based on Lifshitz continuum theory of vdW interactions was developed to calculate the principal many body interactions between arbitrary geometries at all separation distances to a high degree of accuracy through Lifshitz's theory. We applied this numerical implementation to calculate the many body vdW interactions between spherical particles and surfaces with sinusoidally varying roughness profile and also to non spherical particles (cubes, cylinders, tetrahedron etc.) orientated differently with respect to the surface. Our calculations revealed that increasing the surface roughness amplitude decreases the adhesion force and non spherical particles adhere to the surfaces more strongly when their flatter sides are oriented towards the surface. Such practical shapes and structures of particle-surface systems has not been previously considered in resuspension models and this rigorous treatment of vdW interactions provide more realistic adhesion forces between the particle and the surface which can then be coupled with CFD models to improve the predictive capabilities of particle resuspension dynamics.

**Figure 4:**
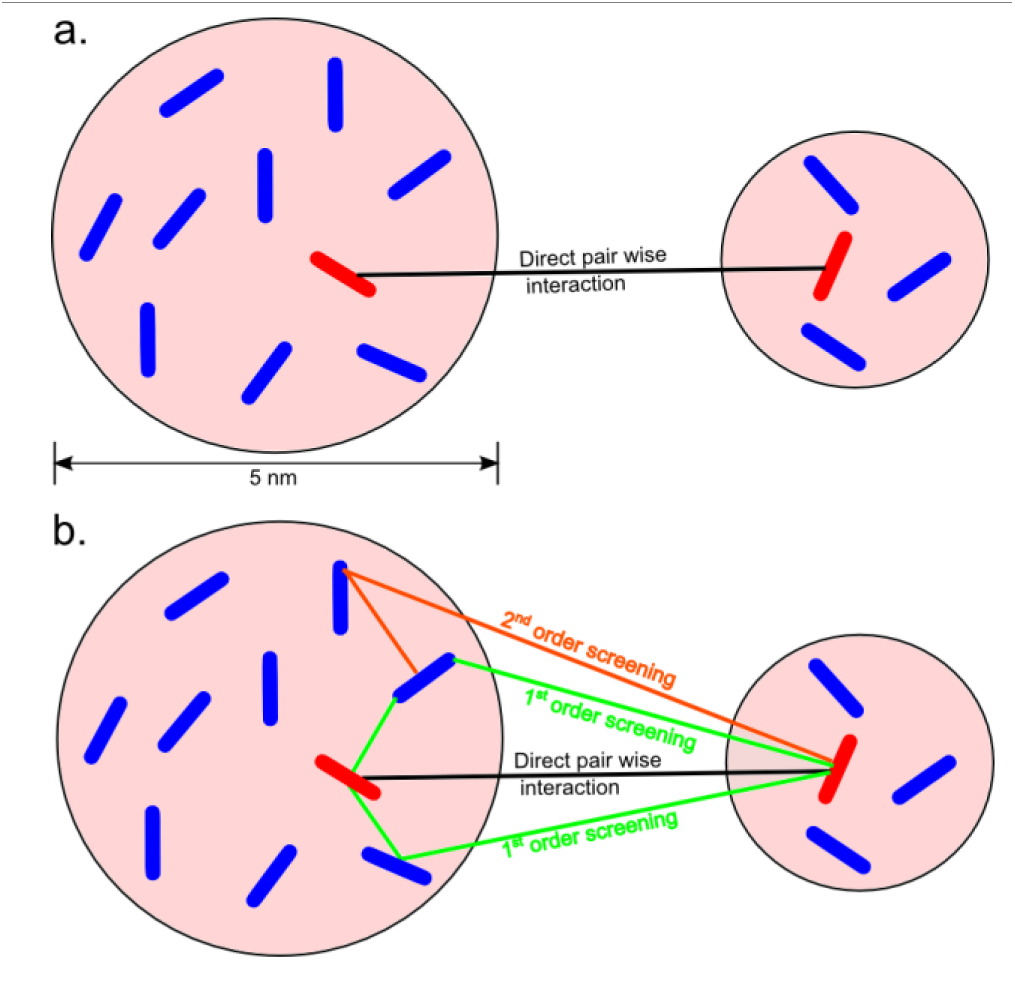
Hamaker two body and Lifshitz continuum interactions. (a) Hamaker's pair wise summation technique where the presence of other molecules does not affect paired interactions. (b) Lifshitz many body interaction takes into account both the direct and indirect (screened) interactions providing a more accurate calculation procedure.

## CONCLUSION

Our analysis of convective flows was also leveraged to create an innovative educational experiences that excite and empower students by helping them recognize how interdisciplinary knowledge can be applied to develop new products and technologies that benefit society. We created novel hands-on activities that introduce chemical engineering students to molecular biology by challenging them to harness microscale natural convection phenomena to perform DNA replication via the PCR [2, 14]. A cognitive assessment reveals that these activities strongly impact student learning in a positive way.

